# Variations in the phagosomal environment of human neutrophils and mononuclear phagocyte subsets

**DOI:** 10.1101/394619

**Authors:** Juliet R. Foote, Amit A. Patel, Simon Yona, Anthony W. Segal

**Author notes:** Corresponding author S.Y. and A.W.S. Joint first author (Juliet R. Foote, Amit A. Patel).

## Abstract

The phagosomal microenvironment has a major influence on the enzyme activity and biology within these organelles. Here we directly compared the phagosomal pH of human neutrophils, monocytes, dendritic cell (DC) and monocyte-derived cells. An unexpected observation was the striking difference in phagosomal environment between the three monocytes subsets. Classical monocytes and neutrophils had alkaline phagosomes, yet non-classical monocytes had more acid phagosomes. Intermediate monocytes had an intermediate phenotype. We next investigated the differences between primary DCs versus *in vitro* monocyte-derived DCs and established that both these cells had acidic phagosomal environments. We also confirmed reports of an alkaline phagosome in “M1” macrophages, and an acidic one in “M2” macrophages. Across all phagocytes, alkalinisation was dependent upon the activity of the NADPH oxidase, as when it was absent in cells from a patient with chronic granulomatous disease (CGD) or was abolished by an inhibitor of the oxidase, diphenyleneiodonium (DPI). An increased alkalinity in the phagosome was associated with more oxidase activity. These data highlight the heterogenous nature of phagocytic vacuoles within the family of mononuclear phagocytes that will dictate the function of these compartments.

**Key points:**

- Phagosomal function depends upon the action of enzymes released into them from cytoplasmic granules.
- The substantial differences in the phagosomal pH in the different phagocytes will affect their compliment of enzymes and their functions.

## Introduction

The ability to internalise particles is an evolutionary conserved process from unicellular organisms to mammalian cells. The downstream purpose of this process varies widely from a simple feeding mechanism to a component of host defence. Within the professional phagocyte family of cells neutrophils, macrophages, monocytes and monocyte-derived cells predominantly kill and digest microbes, whereas others, such as dendritic cells (DCs) are principally involved in antigen presentation. Because all these cells are phagocytic and because their functions are largely conducted within the phagocytic vacuole, it is important to understand the conditions pertaining within that compartment so that they can be related to cellular function.

It has been demonstrated in neutrophils^1,2^ that the phagosomal pH is elevated to about 8.5-9 for at least 30 minutes after phagocytosis. This alkalinisation together with the influx of potassium ions^3^ activates the neutral proteases released from the cytoplasmic granules to kill and digest the ingested microbes. The alkalinisation in neutrophil phagosomes is accomplished by the activity of the NADPH oxidase, NOX2. Superoxide is transported by NOX2 into the phagosome where it forms products such as H_2_O_2_ and H_2_0, consuming protons^1^.

This NOX2 electron transport chain is also present in monocytes, macrophages and DCs^4,5^. Previous studies have demonstrated that NADPH oxidase elevates the pH of phagosomes in monocyte-derived cells stimulated with LPS and IFN-γ termed “M1” macrophages^6^, and has also been shown to be important for antigen processing by DCs^7^ – although questions remain as to how this is accomplished^8^.

Blood monocytes represent a versatile population of cells, composed of several subsets which differ in morphology and transcriptional profiles, described by their location in the blood^9–12^. These monocyte subsets can be distinguished by the membrane expression of CD14 and CD16 in humans^13^ into CD14^+^ CD16^−^ (Classical) monocytes, CD14^+^ CD16^+^ (Intermediate) and CD14^lo^ CD16^+^ (Non-classical) monocytes^14,15^. Monocytes make up around 10% of circulating white blood cells, where classical monocytes up approximately 85% of all monocytes, with intermediate and non-classical monocytes making up the remainder. Classical monocytes are rapidly recruited to sites of infection^16,17^. Non-classical monocytes have been termed patrolling monocytes as they are continuously examining endothelial cell integrity^18^.

DCs bridge the gap between innate and adaptive immunity allowing the immune system to mount precise targeted responses. A key function of the DC is to sense foreign antigen in tissues and present it to T cells, initiating the adaptive immune response^19^. DCs consist of a heterogeneous family of cells that can be classified into plasmacytoid DCs (pDC) or conventional DCs (cDC). cDCs can be further divided into cDC1 or cDC2^20^. cDC1 cells are known to efficiently prime CD8^+^ T cells via cross presentation^21,22^, while, cDC2 have a broad spectrum of activity and can polarize T cells towards Th1, Th2 and Th17 development depending on the antigen presented^23^.

Phagosomal function depends upon the release of the contents of the cytoplasmic granules into this compartment. These are largely enzymes whose optimal pH will vary widely, and so differences in pH in this compartment are likely to have a major influence upon cellular function.

Accordingly, we have undertaken a study to examine this parameter in the different subtypes of mononuclear phagocytes and neutrophils. We have also measured NADPH oxidase activity, as this in turn regulates the pH. Major differences in these parameters in the different cell types examined were observed and are described below.

## Materials and Methods

### Ethics approval

This patient study was carried out in accordance with the recommendations of the Joint UCL/UCLH Committees on the Ethics of Human Research with written informed consent from all subjects. All subjects gave written informed consent in accordance with the Declaration of Helsinki. The protocol was approved by the Joint UCL/UCLH Committees on the Ethics of Human Research (Project numbers 02/0324 and 10/H0806/115). The CGD patient has a mutation in the CYBB gene: c.517delC, predicting p.Leu173CysfsX16.

### Experimental buffer

Balanced salt solution (BSS) buffer contained 156 mM NaCl, 3.0 mM KCl, 1.25mM KH_2_PO_4_, 2 mM MgSO_4_, 2 mM CaCl_2_, 10 mM glucose, 10mM Hepes at pH 7.4.

### Cell isolation and cell culture

Isolation of neutrophils: Human neutrophils from peripheral blood were isolated by dextran sedimentation, centrifugation through Lymphoprep™ (Axis Shield), and hypotonic lysis to remove erythrocytes. Lymphoprep™ or Ficoll™ were used as density gradient mediums, neither has been shown to have any differential effect on cell preparations^24^.

Isolation of monocytes: Monocytes were separated from the interphase layer of whole blood when passed through a density gradient medium (Ficoll™, GE Healthcare), then separated into three populations using flow cytometry cell sorting as described by Patel and colleagues^25^: classical (CD14^+^high, CD16^−^low), non-classical (CD14^−^low, CD16^+^high), and intermediate (CD14^+^, CD16^+^). Cells were resuspended in BSS buffer, then 50-100,000 cell/well in 200 μl volume were added into a poly L-lysine coated Ibidi μ-Slide 15 well angiogenesis plate (Ibidi, Germany) to form an adherent cell monolayer.

Generation of monocyte-derived macrophages: Macrophages were polarised from monocytes, isolated as described above, by method described by Canton and co-workers^6^, in brief: between 8 and 9 x 10^5^ monocytes/well were cultured in an Ibidi μ-Slide 8 well plate (Ibidi, Germany) in RPMI 1640 with 10% fetal bovine serum, 500 U/ml antibiotics (penicillin and streptomycin, ThermoFisher) and 10 mM Hepes buffer (Sigma). For the generation of M1 monocyte-derived cells, the culture medium was supplemented with 60 ng/ml GM-CSF for 5 days, then for a further 2 days with 500 ng/ml LPS and 60 ng/ml IFN-γ. For M2 monocyte-derived cells, 60 ng/ml M-CSF was added for the first 5 days, then 60 ng/ml IL-4 for the final 2 days.

Generation of monocyte-derived dendritic cells: monocytes separated from the interphase layer of whole blood on Ficoll™ were further processed with the Human monocyte enrichment kit (Easy Sep) to isolate classical monocytes. 6 x10^6^ monocytes were cultured for 7 days in 10 cm dishes with 150 ng/ml GM-CSF and 75 ng/ml IL-4 in complete RPMI medium to generate MoDCs^26^.

### Phagosomal pH measurements

The following protocol is described in more detail elsewhere^27^. The wells were washed twice with BSS buffer to remove non-adherent cells, then buffer containing 1 μg/ μl carboxy SNARF-1, AM ester acetate (ThermoFisher) was added for 25 minutes to label the cytosol, and then washed off with BSS buffer. The microscopy plate was mounted on a 37°C heated stage for 15 minutes for acclimatisation before adding approximately 1×10^6^ heat-killed *Candida albicans* (grown from vitroids, Sigma) labelled with SNARF-1 carboxylic acid acetate succinimidyl ester (ThermoFisher) and opsonised with human serum IgG (Vivaglobin). The cells were monitored using a 63× oil immersion on a Zeiss 700 confocal microscope. A snapshot was taken once a minute for 30 minutes when the cells were excited at 555 nm and the emission measured at 560–600 nm and 600-610 nm.

The vacuolar pH was measured using a custom macro within the imaging software ImageJ^28^. At least 20 cells were analysed for each condition within one experiment, *n*=3 unless otherwise stated. The SNARF fluorescence ratio values were converted to pH using the standard curves as described by Levine *et al*^1^: the fluorescence ratios of extracellular SNARF-labelled *Candida* were measured in different buffer solutions (100mM KCl with 50mM buffer salt) from pH 3-13 to construct two standard curves; the fluorescence ratios of SNARF-labelled *Candida* engulfed by human neutrophils were measured after the phagocytosing cells were then subjected to the same buffers with 0.3% saponin; cytoplasmic pH was measured in human neutrophils in the same buffers with nigericin^29^.

### Measurement of phagocytosis

At the end of the kinetic phagosomal pH experiments, trypan blue was added to the cells to quench extracellular Candida fluorescence. Z stacks (8 or 9 1-2-μm sections) were taken in two different random areas of the well for each condition in each experiment, and the total number of cells and total number of cells with at least one engulfed particle were counted using built-in microscope software (Zen, Zeiss).

### Amplex UltraRed assay

The assay was carried out as described^30^ with 50,000 cells in each well. In brief: 50 IU/ml of horseradish peroxidase (Sigma) and 6 μM Amplex ultrared reagent (ThermoFisher) was added to the medium. A basal reading of 3 cycles was recorded before the cells were stimulated by pump injection with 3 μM PMA (Sigma). The fluorescence produced by the oxidation of the Amplex by hydrogen peroxide was measured 30 seconds for 50 minutes at 590 nm after excitation at 540 nm in an Omega FluoStar plate reader (BMG Labtech).

### Statistics

Unless otherwise stated, all experiments were repeated three times. Each graph shows the mean with standard error. Statistical significance was calculated using one-way ANOVA analyses with Bonferroni’s multiple comparison test using the GraphPad Prism 7 software.

## Results

Neutrophils and monocytes have a similar ability to phagocytose pathogens *in vivo*^31^; here we tested if they also have similar downstream phagosomal environments. Neutrophils isolated via dextran sedimentation and density gradient separation yielded a purity of 94% (**Fig. 1a**). Similarly, monocytes were isolated using negative selection resulting in a 91.8% purity. In addition to flow cytometry, the morphology of these cells was assessed **(Fig. 1a and b**). In addition, 68% (SEM ± 4.3%) of neutrophils and 48% (± 4.2%) of monocytes phagocytosed the SNARF-labelled *Candida* but the difference was not significant (data not shown). Neutrophil phagosomes were alkalinised to a pH of approximately 8.5 which was maintained for up to 30 minutes. Similarly, the phagosomes of monocytes also became alkaline, although to a lesser extent reaching and maintaining a pH of about 7.7 (**Fig. 1c**).

**Figure 1.**
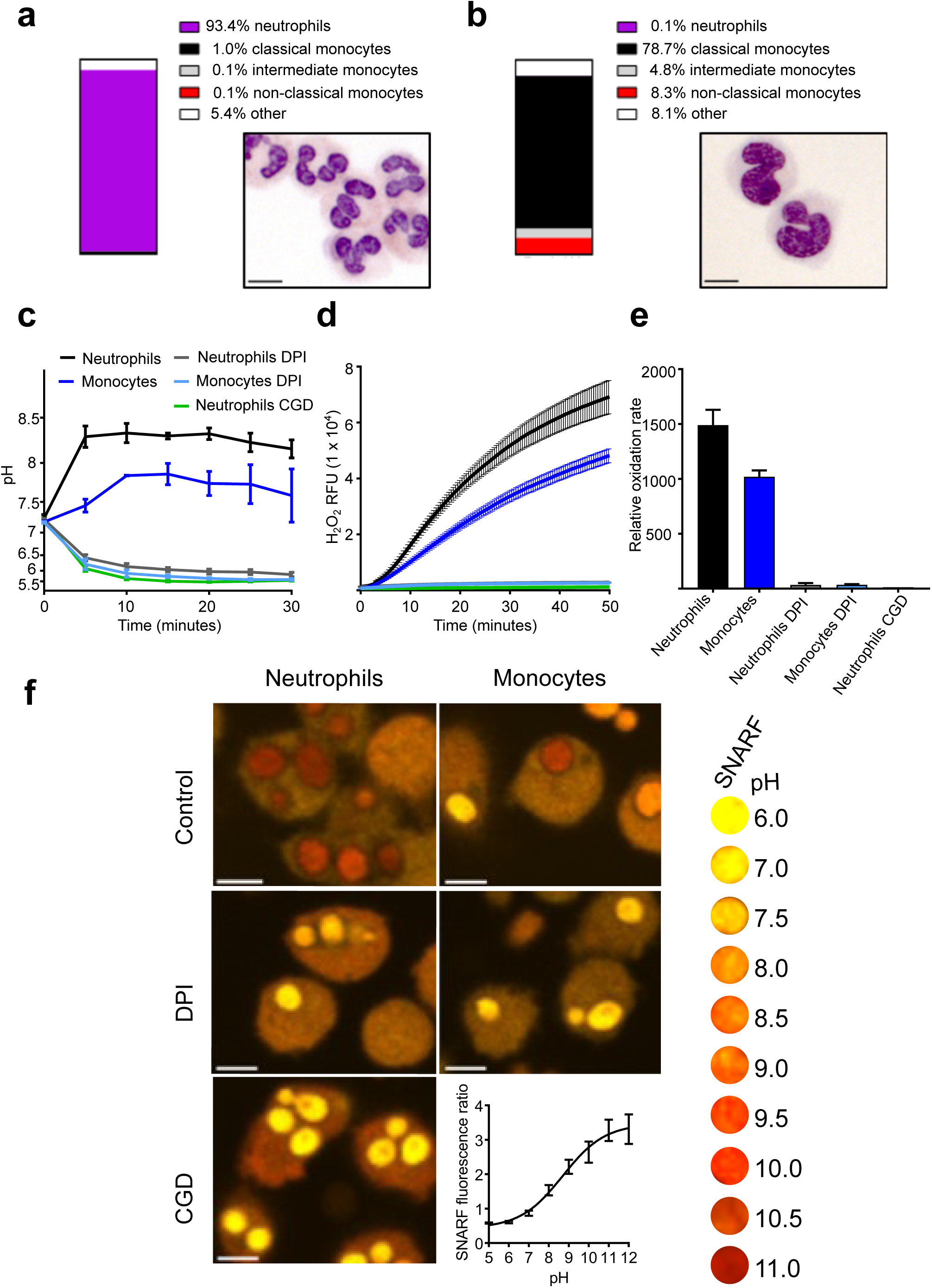
Phagosomal pH and NOX2 activity in human neutrophils and monocytes. Purity and Wright-Giemsa stained preparations of isolated human neutrophils (a) and monocytes (b). Isolated populations were challenged with SNARF-labelled *Candida*. Time course showing the change in phagosomal pH (c), oxidase activity (d) and maximal respiratory rate (e) in neutrophils and monocytes over the first 30 minutes following phagocytosis. These parameters were also measured in a patient with X-linked CGD (*n*=1), and in cells from healthy control treated with DPI (*n*=3), an inhibitor of NOX2. Illustrative images showing *Candida* labelled with the pH indicator SNARF, within neutrophils and monocytes (f) together with a standard curve of SNARF fluorescence against pH (bottom right) and a representative colour key (far right). Calculated p values for pH and respiratory burst between control and DPI and control and CGD for both cell types was <0.001, between DPI and CGD for both cell types was >0.05. Scale bars = 10 μm.

Previous studies have demonstrated that inhibition of NOX2 activity caused phagosomal acidification^32^. To assess whether NOX2 was implicated behind the change in phagosomal pH, cells were treated as above in the presence of the oxidase inhibitor, diphenyleneiodonium (DPI)^33^. While DPI is known to exhibit non-specific effects, X-linked CGD patients have a specific defect in NOX2 which we used to verify the effects of DPI. Oxidase activity, measured by Amplex red oxidation was slower in monocytes as previously described^34,35^ (**Fig. 1e**). Taken together, these data highlight the similarity of monocyte and neutrophil phagosomes. However, the question of differences in phagosome pH between circulating monocyte subsets remains to be answered.

To explore if any differences exist in the phagosomal pH and respiration between monocyte subsets we separated them by FACS into the three subsets based on their CD14 and CD16 surface expression (**Fig. 2a**). Phagocytic capacity of the opsonised heat-killed *Candida* by the subsets was equal (around 50%) in all subsets (data not shown). However, we found that the phagosomal pH in the three subsets was distinct (**Fig. 2b**). The phagosomes of classical monocytes alkalinised to around pH 8.5 by 10 minutes, after which the pH gradually fell to about 7.7. While, non-classical monocytes showed a brief alkalinisation at about 5 minutes after which the pH fell to below 7.0 and remained slightly acidic. The pH of intermediate monocyte phagosomes was intermediate between the other two subsets.

**Figure 2.**
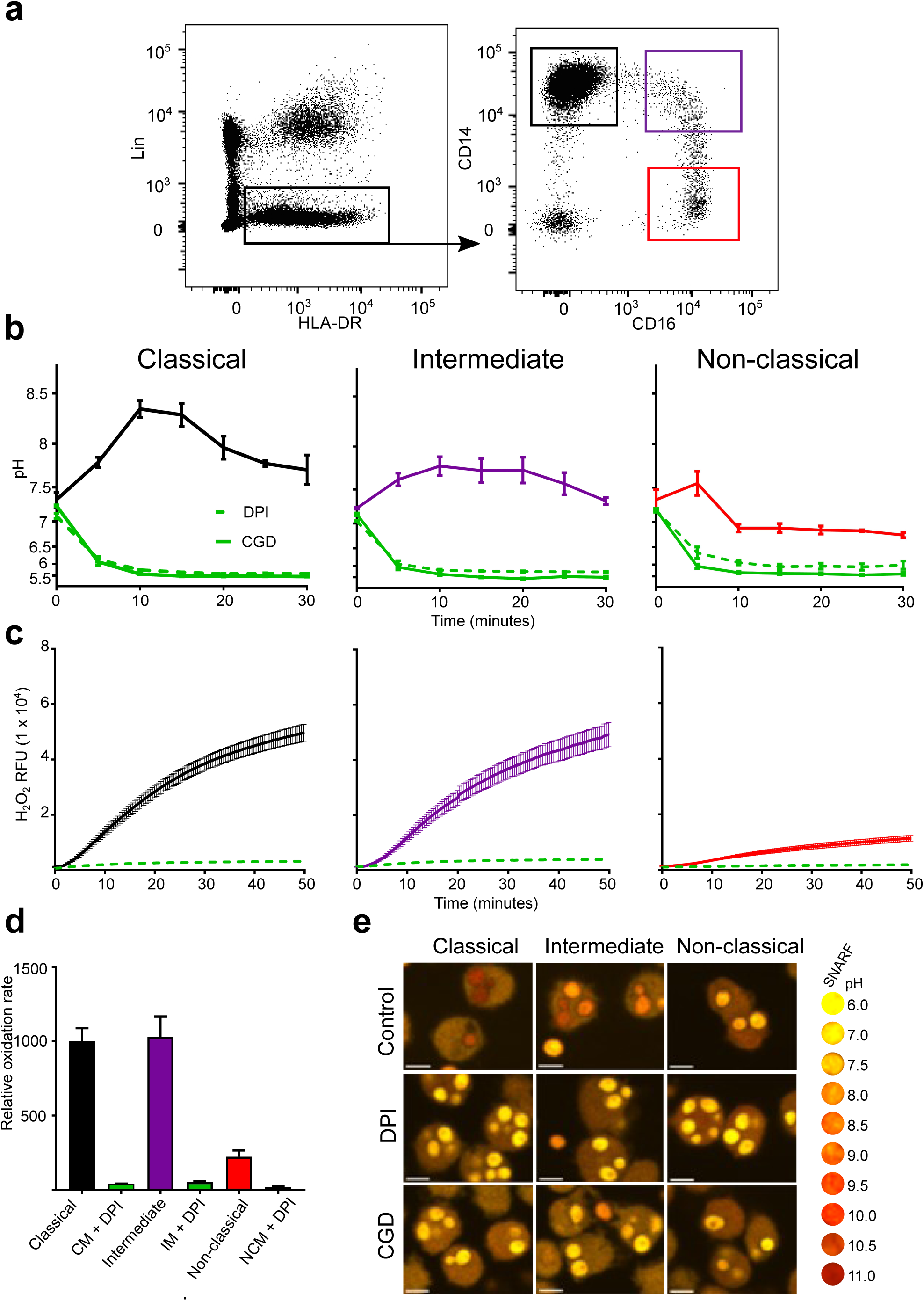
Phagosomal pH and NOX2 activity differed between monocyte subsets. (a) FACS isolation gating strategy of the three monocyte subsets based on CD14 and CD16 expression. (b) Time course of changes in phagosomal pH in monocyte subsets over the first 30 minutes after phagocytosis. The effects of DPI and measurements from an X-linked CGD patient cells are also shown. (c) Respiratory burst activity in subsets in response to stimulation with PMA d) Compiled results of the maximum rates from (c). (e) Representative confocal images of phagocytosed SNARF-*Candida* at 15 minutes post phagocytosis by normal cells, X-linked CGD cells, and healthy cells in the presence of 5 μM DPI. Calculated p values between control conditions for each subset for pH was <0.001, and also for control vs DPI and CGD. No statistical significance between DPI and CGD for all subsets. For the respiratory burst, no significance was found between classical and intermediate controls, but p<0.001 for classical and intermediate controls vs DPI, and p<0.01 for non-classical control vs DPI. Scale bars = 10 μm.

As in neutrophils, the alkalinisation of the phagosomes was produced through the action of NOX2 since the vacuolar pH of all monocyte subsets was acidic in CGD cells or after treatment with DPI. The rate of the respiratory burst was similar for the classical and intermediate monocytes while slower in non-classical monocytes. In all subsets, respiration was significantly inhibited by DPI.

At least three subsets of DCs can isolated from human peripheral blood^36,37^, plasmacytoid DCs (pDC), cDC1 (CD141^+^) and cDC2 (CD1c^+^) (**Fig. 3a**). We could not detect phagocytosis of the SNARF labelled *Candida* in pDCs, and we did not obtain sufficient cDC1s for their adequate examination. In addition, we examined *in vitro* monocyte-derived DCs (**Fig. 3b**) believed to mimic monocytes that differentiate into dendritic-like cells when entering different tissues. It was important to compare primary cDCs with *in vitro* monocyte-derived DC as these cells have been an invaluable tool to many research groups in lieu of primary cDCs (due to their relative scarcity). While it is important to note, the ontogeny^38,39^ and transcriptome^40^ are distinct between naturally occurring cDCs and monocyte-derived DCs.

**Figure 3.**
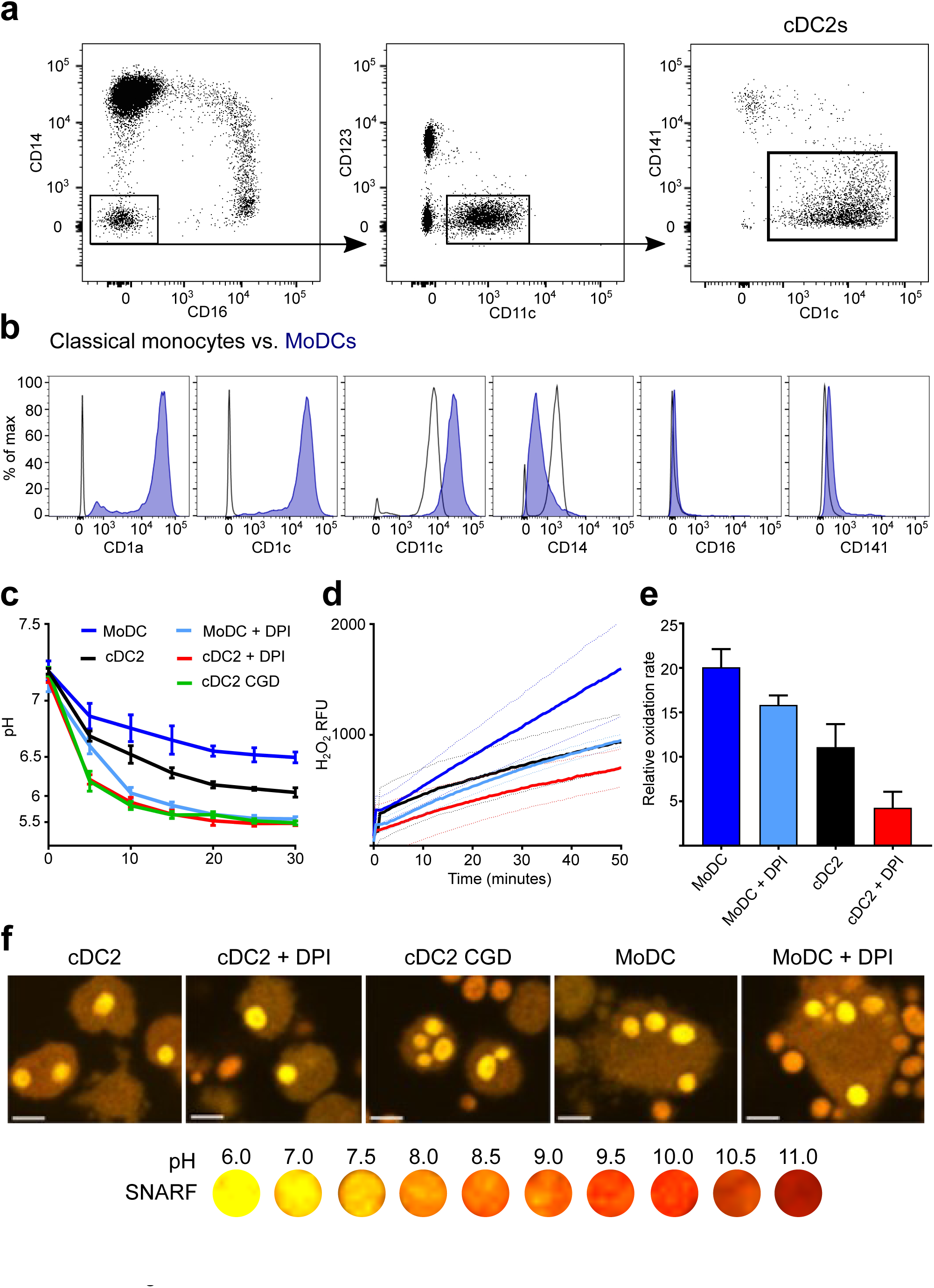
Comparison of blood CD1c^+^ DCs and monocyte-derived DCs. (a) Gating strategy used to isolate CD1c^+^ (cDC2s) using FACS; (b) *in vitro* derived-MoDCs were phenotypically validated according to surface markers by comparison with their precursor classical monocytes ^58,59^, a representative plot is shown based on three experiments; (c) Time course of changes in the pH in the phagosomes of MoDCs and cDC2s, from a healthy subject, with and without DPI, and cDC2s from a CGD patient; d) NADPH oxidase activity in response to PMA stimulation in moDCs and cDC2s in the presence and absence of DPI; (e) maximal rates of respiration from (d); (f) Representative confocal images of each subtype 15 minutes after phagocytosis. Calculated p values for phagosomal pH time courses: p<0.05 for cDC2s vs MoDCs <0.05, p<0.001 for cDC2s vs DPI and for cDC2s vs CGD, p<0.001 for MoDCs vs DPI. For the respiratory burst, p<0.01 for cDC2s vs MoDCs, no significance was found between cDC2s and cells with DPI, but p<0.001 for MoDCs vs DPI. Scale bar = 10 μm.

We found that the phagosomal pH was slightly less acidic in MoDCs compared to primary circulating cDC2s (**Fig. 3c**). Upon the addition of DPI, the pH decreased further in both cell types (p<0.001 in both cell types), resembling levels observed CGD patient derived cDC2s. Interestingly, cDC2s were much less phagocytic than MoDCs, with a mean percentage of phagocytosing cells 23.4 ± 4.6 SEM and 60.0 ± 3.1 respectively, which may correspond with previous findings that MoDCs are superior at receptor mediated endocytosis of immune complexes^26^.

The amount and rate of hydrogen peroxide production in response to PMA was much lower in both types of DCs (**Fig. 3d and e**) in comparison to neutrophils and classical monocytes **(Fig. 1d and e**). However, MoDCs were found to produce more H2O2 than cDC2s (p<0.01). The respiratory burst of MoDCs was further lowered by DPI (p<0.001). The change in oxidase activity of cDC2 cells was not significantly affected by DPI.

We next measured phagosomal pH monocyte-derived “M1” and “M2” cells and the undifferentiated state, “M0” cells. We first confirmed that the identity of the subsets as shown in **Figure 4a**. The phagosomal pH of classically activated monocyte-derived cells termed “M1” macrophages, were alkaline at 5 minutes after phagocytosis, similar to neutrophils, and maintained a phagosomal pH of ∼8.5 (**Fig. 4b**). In contrast, alternatively activated monocyte-derived cells “M2” macrophages acidified their phagosomes in a similar fashion to non-classical monocytes. M0 reached a level between that of M1 and M2 macrophages, this was due to both acidic and alkaline phagosomes (**Fig. 4c**). There was no significant change in pH between “M2” cells with and without DPI, suggesting that NOX2 does not regulate the phagosomal pH in these cells. These results corroborate previous studies by Canton and colleagues^6^.

**Figure 4.**
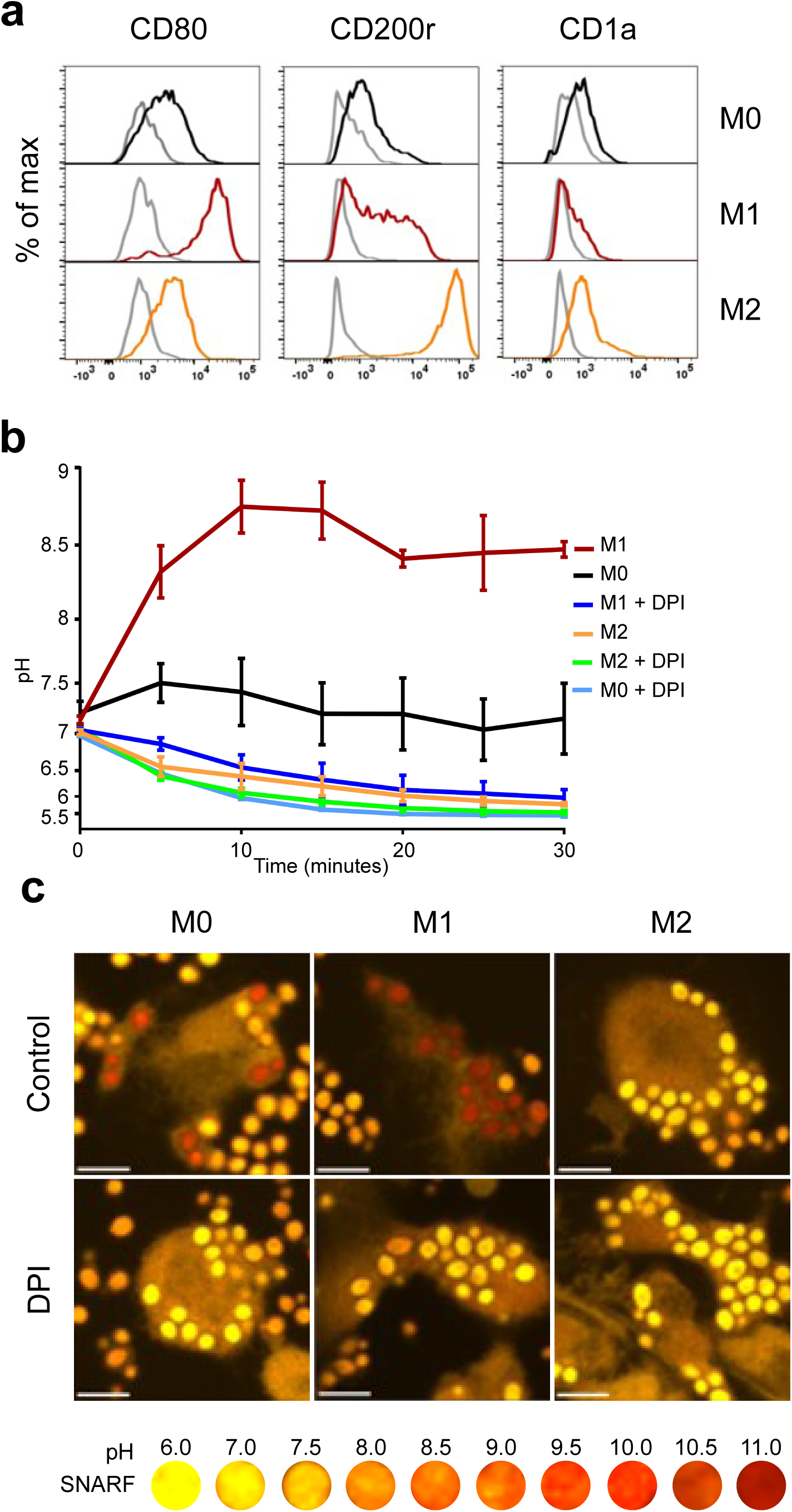
Polarised macrophages have different phagosomal pH profiles. (a) Phenotypic validation of monocyte-derived macrophage differentiation from undifferentiated macrophages (MO) into M1 and M2 macrophages using flow cytometry. CD80 was used as a marker for M1 differentiation, CD200r for M2, and CD1a to exclude dendritic cells differentiation. (b) Time course of phagosomal pH in the three monocyte-derived cell populations. (c) Representative confocal images of each macrophage subtype 15 minutes after phagocytosis. Calculated p values for phagosomal pH time courses: p<0.001 for between all macrophage controls, P<0.001 for M1 vs M1 DPI and M0 vs M0 DPI, ns for M2 vs M2 DPI. Scale bar = 10 μm.

## Discussion

It is currently thought that the role of the phagosome in professional phagocytes falls into two main categories, pathogen killing and digestion, or antigen processing and presentation. Both these roles require the activity of digestive enzymes and these in turn are highly dependent upon the conditions within the phagocytic vacuole. Previously, the pH of human monocyte vacuoles has been described as incredibly acidic. Jacques and Bainton used visible dyes (limited to detection ranges below pH 7) to show that ∼40% of monocytes had acidified to a pH of 4.5 to 5.0 within 10 minutes^41^ whilst others used fluorescein fluorescence to determine the pH at about 5-6 ^42^. There appeared to be a discrepancy between these low pHs and the fact that monocytes have an active respiratory burst^43^, which has been shown in neutrophils to elevate the pH within the phagocytic vacuole^1,2^. In addition, it is known that the neutrophil and monocyte cytoplasmic granules contain a variety of enzymes, including lysozyme and myeloperoxidase^44–46^, although more poorly classified in monocytes. These enzymes vary in pH optima, for example myeloperoxidase tested in neutrophils is pH 5-6^1^ whereas lysozyme (tested in serum) lies between pH 8-9^47^. Together with our observation of difference in phagosomal pH in monocyte subsets, this may indicate that (a) these enzymes are present in different subsets of monocytes, (b) they are contained within different granules which degranulate at different times in the evolution of the vacuole, or (c) that the timing of their activity varies with temporal changes in the vacuole.

Our results show that like neutrophils, classical monocytes develop alkaline vacuoles. In addition, the duration and extent of vacuolar alkalinisation varied considerably between monocyte subsets. This suggests functional diversity exists between monocyte subsets.

The results on polarised macrophages mirrored those described by Grinstein^48^, however we included the observation that undifferentiated monocyte derived cells (without incubation with cytokines) were more acidic than the “M1” cells yet distinct from the “M2” monocyte-derived cells. The phagosomal pHs of “M1” and “M0” macrophages were acidified significantly with the addition of DPI, whereas “M2” macrophages were unaffected. It is apparent that different mechanisms involving NOX2 and regulation of the pH are being switched on in these phenotypes and offers a model to investigate them further.

The role of the NADPH oxidase on antigen handling by dendritic cells is a point of contention in the field. A number of groups report that mouse bone marrow derived DCs^49,50^ and human blood DCs^51^ have an alkaline phagosomal pH (pH 7-8), which is elevated by the NADPH oxidase, and is necessary for optimal antigen processing and presentation, as the lysosomal enzymes are activated by an acidic environment and degrade the antigen^52,53^.

On the other hand, Rybicka and colleagues^8^ reported that mouse BMDCs and splenic DCs had acidic phagosomal pHs that were unaltered by the activity of the oxidase, and proposed that it was the reducing environment, linked to the pH, rather than the proton concentration, that was important for correct antigen presentation. Another group came to the same conclusions also using mouse BMDCs^54^, while Westcott found that *Listeria*-containing mouse BMDC phagosomes became acidic within the first hour after ingestion^55^. In human MoDCs, a group reported phagosomal pH of just under pH 7 maintained 90 minutes after phagocytosis^56^, while another found phagosomes to be more acidic at around pH 5^57^ but 4 hours after particle ingestion.

In accordance with these findings, we did not observe the vacuolar pH of either cDC2s or the majority of MoDCs to become alkaline at any stage. However, we agreed with Savina^53^ and colleagues that the oxidase does influence the vacuolar pH which we found to be lower when it was absent in CGD or inhibited by DPI.

The fact that we found the vacuoles of DCs to be acidic whereas others have described them as alkaline could reflect species differences, the majority of these studies have been conducted on mouse cells, or in differences in the handling of the cells. Like macrophages, most researchers differentiate cells in culture with differing combinations and concentrations of growth factors and stimuli, either from bone marrow (BMDCs, mainly in mice) or from circulating blood monocytes (MoDCs, mainly from humans).

To date characterisation of these blood derived mononuclear cells has been directed to their differentiation in terms of their surface markers, lineages, and immunological functions. The results of this study indicate that the subsets of these mononuclear cells maintain very different physical environments within their vacuoles which will have major effects on the enzymology within these compartments. To gain a clearer understanding of macrophage heterogeneity, primary macrophage phagosomes from distinct tissue i.e. alveolar macrophages or Langerhans cells could be examined. Future work into the identification of the enzymes released into the vacuoles of these different cellular subtypes, and their differential functions, will enable a better understanding of the function of these cells.

## Acknowledgements

J.R.F. was supported on an Irwin Joffe Memorial Fellowship. A.A.P. was supported by an EPSRC studentship. We thank Siobhan Burns for clinical assistance, Dirk Roos for sequencing data of the CGD patient, Jamie Evans for flow cytometry assistance. Finally, we thank all the volunteers and patients for donating their blood.

## Authorship

Contribution: J.R.F. and A.A.P. isolated and cultured cells and performed flow cytometry experiments, A.A.P. performed purity analyses, J.R.F. performed and analysed microscopy experiments and Amplex UltraRed assays and made the figures. S.Y. and A.W.S. designed the research, all authors wrote the paper.

Conflict-of-interest disclosure: The authors declare no competing financial interests.

Correspondence: Simon Yona and Anthony Segal, Division of Medicine, University College London, U.K.;

## References

1. Levine AP, Duchen MR, de Villiers S, Rich PR, Segal AW. Alkalinity of neutrophil phagocytic vacuoles is modulated by HVCN1 and has consequences for myeloperoxidase activity. PLoS One. 2015;10(4):e0125906.

2. Segal AW, Geisow M, Garcia R, Harper A, Miller R. The respiratory burst of phagocytic cells is associated with a rise in vacuolar pH. Nature. 1981;290(5805):406–409.

3. Reeves EP, Lu H, Jacobs HL, et al. Killing activity of neutrophils is mediated through activation of proteases by K+ flux. Nature. 2002;416(6878):291–7.

4. Segal AW, Garcia R, Goldstone H, Cross AR, Jones OT. Cytochrome b-245 of neutrophils is also present in human monocytes, macrophages and eosinophils. Biochem. J. 1981;196(1):363–7.

5. Elsen S, Doussière J, Villiers CL, et al. Cryptic O2- -generating NADPH oxidase in dendritic cells. J. Cell Sci. 2004;117(Pt 11):2215–26.

6. Canton J, Khezri R, Glogauer M, Grinstein S. Contrasting phagosome pH regulation and maturation in human M1 and M2 macrophages. Mol. Biol. Cell. 2014;25(21):3330–41.

7. Mantegazza AR, Savina A, Vermeulen M, et al. NADPH oxidase controls phagosomal pH and antigen cross-presentation in human dendritic cells. Blood. 2008;112(12):.

8. Rybicka JM, Balce DR, Chaudhuri S, Allan ERO, Yates RM. Phagosomal proteolysis in dendritic cells is modulated by NADPH oxidase in a pH-independent manner. EMBO J. 2012;31(4):932–44.

9. Geissmann F, Jung S, Littman DR. Blood Monocytes Consist of Two Principal Subsets with Distinct Migratory Properties. Immunity. 2003;19(1):71–82.

10. Ingersoll MA, Spanbroek R, Lottaz C, et al. Comparison of gene expression profiles between human and mouse monocyte subsets. Blood. 2010;115(3):e10–9.

11. Yona S, Jung S. Monocytes: subsets, origins, fates and functions. Curr. Opin. Hematol. 2010;17(1):53–59.

12. Mildner A, Schönheit J, Giladi A, et al. Genomic Characterization of Murine Monocytes Reveals C/EBPβ Transcription Factor Dependence of Ly6C−Cells. Immunity. 2017;46(5):849–862.e7.

13. Ziegler-Heitbrock L, Ancuta P, Crowe S, et al. Nomenclature of monocytes and dendritic cells in blood. Blood. 2010;116(16):e74–80.

14. Passlick B, Flieger D, Ziegler-Heitbrock H. Identification and characterization of a novel monocyte subpopulation in human peripheral blood. Blood. 1989;74(7):.

15. Wong KL, Tai JJ-Y, Wong W-C, et al. Gene expression profiling reveals the defining features of the classical, intermediate, and nonclassical human monocyte subsets. Blood. 2011;118(5):e16–31.

16. Liao C-T, Andrews R, Wallace LE, et al. Peritoneal macrophage heterogeneity is associated with different peritoneal dialysis outcomes. Kidney Int. 2017;91(5):1088–1103.

17. Serbina N V, Pamer EG. Monocyte emigration from bone marrow during bacterial infection requires signals mediated by chemokine receptor CCR2. Nat. Immunol. 2006;7(3):311–317.

18. Auffray C, Fogg D, Garfa M, et al. Monitoring of blood vessels and tissues by a population of monocytes with patrolling behavior. Science. 2007;317(5838):666–70.

19. Merad M, Sathe P, Helft J, Miller J, Mortha A. The dendritic cell lineage: ontogeny and function of dendritic cells and their subsets in the steady state and the inflamed setting. Annu. Rev. Immunol. 2013;31:563–604.

20. Guilliams M, Ginhoux F, Jakubzick C, et al. Dendritic cells, monocytes and macrophages: a unified nomenclature based on ontogeny. Nat. Rev. Immunol. 2014;14(8):571–578.

21. Haniffa M, Collin M, Ginhoux F. Ontogeny and Functional Specialization of Dendritic Cells in Human and Mouse. Adv. Immunol. 2013;120:1–49.

22. Imai T, Kato Y, Kajiwara C, et al. Heat shock protein 90 (HSP90) contributes to cytosolic translocation of extracellular antigen for cross-presentation by dendritic cells. Proc. Natl. Acad. Sci. U. S. A. 2011;108(39):16363–8.

23. Schlitzer A, McGovern N, Teo P, et al. IRF4 Transcription Factor-Dependent CD11b+ Dendritic Cells in Human and Mouse Control Mucosal IL-17 Cytokine Responses. Immunity. 2013;38(5):970–983.

24. Yeo C, Saunders N, Locca D, et al. Ficoll-PaqueTM versus LymphoprepTM: a comparative study of two density gradient media for therapeutic bone marrow mononuclear cell preparations. Regen. Med. 2009;4(5):689–696.

25. Patel AA, Zhang Y, Fullerton JN, et al. The fate and lifespan of human monocyte subsets in steady state and systemic inflammation. J. Exp. Med. 2017;214(7):.

26. Andersson LIM, Cirkic E, Hellman P, Eriksson H. Myeloid blood dendritic cells and monocyte-derived dendritic cells differ in their endocytosing capability. Hum. Immunol. 2012;73(11):1073–1081.

27. Foote JR, Levine AP, Behe P, Duchen MR, Segal AW. Imaging the neutrophil phagosome and cytoplasm using a ratiometric pH indicator. J. Vis. Exp. 2017;2017(122):.

28. Schneider CA, Rasband WS, Eliceiri KW. NIH Image to ImageJ: 25 years of image analysis. Nat. Methods. 2012;9(7):671–675.

29. Morgan D, Capasso M, Musset B, et al. Voltage-gated proton channels maintain pH in human neutrophils during phagocytosis. Proc. Natl. Acad. Sci. U. S. A. 2009;106(42):18022–18027.

30. Maini A, Foote JR, Hayhoe R, et al. Monocyte and Neutrophil Isolation, Migration, and Phagocytosis Assays. Curr. Protoc. Immunol. 2018;122(1):e53.

31. Rydström A, Wick MJ. Infection Salmonella Tissue during Oral Function in the Gut-Associated Lymphoid Monocyte Recruitment, Activation, and. J Immunol Ref. 2018;178:5789–5801.

32. Jankowski A, Grinstein S. Modulation of the cytosolic and phagosomal pH by the NADPH oxidase. Antioxid. Redox Signal. 2002;4(1):61–8.

33. Cross AR, Jones OT. The effect of the inhibitor diphenylene iodonium on the superoxide-generating system of neutrophils. Specific labelling of a component polypeptide of the oxidase. Biochem. J. 1986;237(1):111–6.

34. de Rossi L, Gott K, Horn N, et al. Xenon preserves neutrophil and monocyte function in human whole blood. Can. J. Anesth. 2002;49(9):942–945.

35. Chu J, Song HH, Zarember KA, Mills TA, Gallin JI. Persistence of the bacterial pathogen Granulibacter bethesdensis in chronic granulomatous disease monocytes and macrophages lacking a functional NADPH oxidase. J. Immunol. 2013;191(6):3297–307.

36. See P, Dutertre C-A, Chen J, et al. Mapping the human DC lineage through the integration of high-dimensional techniques. Science. 2017;356(6342):eaag3009.

37. Villani A-C, Satija R, Reynolds G, et al. Single-cell RNA-seq reveals new types of human blood dendritic cells, monocytes, and progenitors. Science. 2017;356(6335):eaah4573.

38. Lee J, Breton G, Oliveira TYK, et al. Restricted dendritic cell and monocyte progenitors in human cord blood and bone marrow. J. Exp. Med. 2015;212(3):385–99.

39. Breton G, Lee J, Zhou YJ, et al. Circulating precursors of human CD1c+ and CD141+ dendritic cells. J. Exp. Med. 2015;212(3):401–13.

40. Robbins SH, Walzer T, Dembélé D, et al. Novel insights into the relationships between dendritic cell subsets in human and mouse revealed by genome-wide expression profiling. Genome Biol. 2008;9(1):R17.

41. Jacques Y V, Bainton DF. Changes in pH within the phagocytic vacuoles of human neutrophils and monocytes. Lab. Invest. 1978;39(3):179–85.

42. Horwitz MA, Maxfield FR. Legionella pneumophila inhibits acidification of its phagosome in human monocytes. J. Cell Biol. 1984;99(6):1936–43.

43. Seres T, Knickelbein RG, Warshaw JB, Johnston RB. The Phagocytosis-Associated Respiratory Burst in Human Monocytes Is Associated with Increased Uptake of Glutathione. J. Immunol. 2000;165(6):3333–3340.

44. Resnitzky P, Shaft D, Yaari A, Nir E. Distinct intracellular lysozyme content in normal granulocytes and monocytes: a quantitative immunoperoxidase and ultrastructural immunogold study. J. Histochem. Cytochem. 1994;42(11):1471–7.

45. Lewis CE, Mccarthy SP, Lorenzen J, Mcgee Nuffield D. Differential effects of LPS, IFN-y and TNFa on the secretion of lysozyme by individual human mononuclear phagocytes: relationship to cell maturity. 1990.

46. Senior RM, Campbell EJ. Cathepsin G in human mononuclear phagocytes: comparisons between monocytes and U937 monocyte-like cells. J. Immunol. 1984;132(5):2547–51.

47. Wardlaw AC. The complement-dependent bacteriolytic activity of normal human serum. I. The effect of pH and ionic strength and the role of lysozyme. J. Exp. Med. 1962;115(6):1231–49.

48. Canton J. Phagosome maturation in polarized macrophages. J. Leukoc. Biol. 2014;96(5):729–38.

49. Liu X, Lu L, Yang Z, et al. The neonatal FcR-mediated presentation of immune-complexed antigen is associated with endosomal and phagosomal pH and antigen stability in macrophages and dendritic cells. J. Immunol. 2011;186(8):4674–86.

50. Dingjan I, Verboogen DR, Paardekooper LM, et al. Lipid peroxidation causes endosomal antigen release for cross-presentation. Sci. Rep. 2016;6(1):22064.

51. Segura E, Durand M, Amigorena S. Similar antigen cross-presentation capacity and phagocytic functions in all freshly isolated human lymphoid organ-resident dendritic cells. J. Exp. Med. 2013;210(5):1035–47.

52. Jancic C, Savina A, Wasmeier C, et al. Rab27a regulates phagosomal pH and NADPH oxidase recruitment to dendritic cell phagosomes. Nat. Cell Biol. 2007;9(4):367–378.

53. Savina A, Jancic C, Hugues S, et al. NOX2 Controls Phagosomal pH to Regulate Antigen Processing during Crosspresentation by Dendritic Cells. Cell. 2006;126(1):205–218.

54. Salao K, Jiang L, Li H, et al. CLIC1 regulates dendritic cell antigen processing and presentation by modulating phagosome acidification and proteolysis. Biol. Open. 2016;5(5):620–30.

55. Westcott MM, Henry CJ, Amis JE, Hiltbold EM. Dendritic cells inhibit the progression of Listeria monocytogenes intracellular infection by retaining bacteria in major histocompatibility complex class II-rich phagosomes and by limiting cytosolic growth. Infect. Immun. 2010;78(7):2956–65.

56. Kourjian G, Rucevic M, Berberich MJ, et al. HIV Protease Inhibitor-Induced Cathepsin Modulation Alters Antigen Processing and Cross-Presentation. J. Immunol. 2016;196(9):3595–607.

57. Thiele L, Merkle HP, Walter E. Phagocytosis and Phagosomal Fate of Surface-Modified Microparticles in Dendritic Cells and Macrophages. 2003.

58. Ohradanova-Repic A, Machacek C, Fischer MB, Stockinger H. Differentiation of human monocytes and derived subsets of macrophages and dendritic cells by the HLDA10 monoclonal antibody panel. Clin. Transl. Immunol. 2016;5(1):e55.

59. Goudot C, Coillard A, Villani AC, et al. Aryl Hydrocarbon Receptor Controls Monocyte Differentiation into Dendritic Cells versus Macrophages. Immunity. 2017;

